# Identifying core biological processes distinguishing human eye tissues with precise systems-level gene expression analyses and weighted correlation networks

**DOI:** 10.1101/136960

**Authors:** John M Bryan, Temesgen D Fufa, Kapil Bharti, Brian P Brooks, Robert B Hufnagel, David M McGaughey

## Abstract

The human eye is built from several specialized tissues which direct, capture, and pre-process information to provide vision. The gene expression of the different eye tissues has been extensively profiled with RNA-seq across numerous studies. Large consortium projects have also used RNA-seq to study gene expression patterning across many different human tissues, minus the eye. There has not been an integrated study of expression patterns from multiple eye tissues compared to other human body tissues. We have collated all publicly available healthy human eye RNA-seq datasets as well as dozens of other tissues. We use this fully integrated dataset to probe the biological processes and pan expression relationships between the cornea, retina, RPE-choroid complex, and the rest of the human tissues with differential expression, clustering, and GO term enrichment tools. We also leverage our large collection of retina and RPE-choroid tissues to build the first human weighted gene correlation networks and use them to highlight known biological pathways and eye gene disease enrichment. We also have integrated publicly available single cell RNA-seq data from mouse retina into our framework for validation and discovery. Finally, we make all these data, analyses, and visualizations available via a powerful interactive web application (https://eyeintegration.nei.nih.gov/).

## Introduction

The human eye is a highly specialized organ using several distinct tissues to focus and capture light and begin processing it into visual information. Light passes through the cornea and the lens which focus the light onto the retina (1). The rod and cone photoreceptors of the retina capture the light and transmits visual information that is processed by a network of retinal synapses and passed through the optic nerve to the brain (2). The retinal pigment epithelium (RPE) is responsible for absorbing scattered light and providing nutrition, maintaining ionic homeostasis, and waste product processing for the photoreceptors, as well as mediating immune function for the retina and eye (3). The RPE and outer neural retina is supported by and connected to the vascular system of the body via the choroid (4).

Many genetic disorders affect the function of the various eye tissues and cause vision perturbation or loss. The genetics of eye diseases range from monogenic Mendelian disorders to complex multi-gene system perturbations that are modified by environmental influences. While at least 316 identified genes underlying retinal diseases have been identified, recent comprehensive next generation sequencing studies fail to find the cause of a variety of inherited retinal diseases like cone-rod dystrophies or retinitis pigmentosa 40–60% of the time (5–7). In an example of complex disease, age-related macular degeneration (AMD), which is believed to be caused by dysfunction of the RPE and choroid, genome-wide association studies (GWAS) have identified dozens of genomic locations associated with the disease. Still it is very difficult to pinpoint the causative gene or genes (8).

An valuable tool in understanding basic biology and unravelling the causes of disease has been the analysis of gene expression profiles. The Genotype-Tissue Expression (GTEx) Project has compiled nearly 10,000 individual tissue human RNA-seq samples and shared the data via a powerful and easy-to-use web portal (9). GTEx data has been used to help filter variants in GWAS studies, to build networks to identify candidate testis cancer genes, to help identify pathogenic mutations in an epilepsy cohort, and to identify a genetic variant linking folate homeostasis to warfarin response (10–14). Notably, the eye was not included as a tissue for this project. Because the vision community has been adopting RNA-seq for profiling different components of the eye, there is a large and growing set of useful transcriptome data. However, each study uses different bioinformatic processes to analyze their transcriptomes and the full genome-wide expression values are difficult to obtain, analyze, and visualize across studies. Therefore, utility of these resources ought to be optimized to similar effect as for other tissues.

We have collated all publicly available human eye tissue RNA-seq data and processed it with a robust and consistent bioinformatics process. We also have brought in a substantial portion of the GTEx project RNA-seq data to provide a comparison set to the eye tissues. Our full data-set holds 1027 samples. This comprehensive and consistently processed pan-eye and human data set allows for several novel analyses: first, to probe the relationships within cornea, retina, and RPE tissues and between eye tissues and other human tissues; second, to look for overarching patterns in gene expression and shared biology in differentially expressed genes between the eye tissues; and, third, we use the large collated retina and RPE samples to build gene correlation networks for both. A single cell mouse RNA-seq retina dataset has also been integrated to validate the retina gene correlation networks.

To maximize utility of this project to all researchers, we have also created a freely available web application that allows quick and powerful access to the expression profiles of nearly 20,000 genes across 177 human eye tissue RNA-seq sets and 853 GTEx tissue RNA-seq sets, the two gene networks, and the 10,000 plus cells of the mouse retina single cell dataset (https://eyeIntegration/nei.nih.gov/).

## Materials and Methods

### Identification of normal human eye RNA-seq data-sets and tissue labelling

The entire SRA dataset was downloaded as a SQL file on January 19^th^, 2017 with the SRAdb R package. The following keywords were used in a partial-matching case-insensitive (e.g. ‘retina’ would match ‘RETINAL’) search: ‘RPE’, ‘macula’, ‘fovea’, ‘retina’, ‘choroid’, ‘sclera’, ‘iris’, ‘lens’, ‘cornea’, and ‘eye.’ These keywords were matched against the following fields in the SRA: ‘study_abstract’, ‘experiment_name’, ‘study_name’, ‘sample_ID’, ‘sample_name’, ‘study_title’, ‘study_description’ in human samples with a ‘library_source’ of ‘transcriptomic’ and filtering out miRNA studies. Study titles, abstracts, and other fields were checked by hand for inclusion in this study for whether they were genuine eye studies of normal (non-disease, non-mutated, no chemical modification) human eye tissue. The SRA metadata for the GTEx project was also pulled by searching for the study accession ‘SRP012682.’ Our script enabling search of the SRA for eye tissues is provided as ‘sraDB_search_select.R’

For reproducibility, the meta-data for each sample was parsed with our script ‘parse_sample_attribute.R’ to label the eye tissue (cornea, lens, eye-lid, retina, RPE, ESC) and its origin (immortalized cell-line, cell-line derived from ESC, fetal tissue, or adult tissue). This script has been written to handle the wide variety of metadata usage by the 21 research projects and the script likely would need to be modified to handle new eye samples. The GTEx tissue were labeled by tissue or sub tissue by parsing the GTEx SRA metadata for ‘histological type’ and ‘body site’, respectively with the ‘parse_sample_attribute.R’ script.

### Efficient quantification of gene expression across 1027 samples

Two studies had their raw RNA-seq data accessioned with dbGaP (9, 15). We obtained access to these studies under dbGaP study #115588. Raw sequence data for these two studies were pulled and converted to fastq with the sratoolkit (2.8.0) fastq-dump tool. The remaining raw fastq data was pulled from NCBI via ftp, with the wget calls created by the script ‘sra_to_fastq.R’. The one exception was the E-MTAB-4377 resource which was only available in the bam format as of January 19^th^ 2017 from European Bioinformatics Institutes ArrayExpress archive (16). The bam files were downloaded, then converted to fastq with the Picard SamToFastq (2.1.1) program (https://broadinstitute.github.io/picard/).

The raw fastq read files were loaded into salmon (0.7.2) with –seqBias and –gcBias flags against the Gencode Release 25 protein-coding transcript sequences fasta file to perform transcript-level quantification (17, 18). The Gencode gene names are used across this study. To improve specificity of the gene expression, transcripts with low abundance across all tissues were removed from the fasta file, and Salmon was re-run as per Soneson et. al (19). The filtered fasta file is provided in the source code as ‘gencode.v25.pc_transcripts.commonTx.fa.gz.’ and the Salmon script as ‘run_salmon.sh.’ To improve sensitivity and specificity, the transcript-level quantifications were merged to the gene-level and the length scaled transcripts per kilobase million (TPM) calculations were done with the R library tximport (1.2.0) (20) in our ‘ calculate_lengthScaledTPM.R’ script.

### Multi-step process to remove samples with low overall gene expression counts, quantile normalize samples by tissue, then cluster to identify outliers

A multi-stage process was then used on the full data set to remove outlier samples (either because of overall low gene expression levels or from clustering with the incorrect tissue group). Genes with zero to extremely low expression across the entire data-set were removed. While we found several mislabeled GTEx samples, this has been noticed before (21). Samples with a median TPM value < 50 were removed as these were outliers in terms of overall gene expression coverage. This step alone removed all of the lens samples, 20 RPE, 15 retina, and 16 ESC samples (Supplementary Materials, Table S1, S2 and Figure S2). To alleviate potential batch effects between the samples from different studies, the TPM values were quantile normalized within tissues and globally simultaneously with the qsmooth algorithm (22) (Supplementary Materials, Figure S7).

Finally, the remaining samples were dimensionality reduced with t-SNE, then clustered with DBSCAN. The performance of t-SNE is sensitive to the perplexity parameter, which weighs local versus global relationships. We found for our study that perplexities ranging from 30-50 performed the most reliably (data not shown). For the all-sample t-SNE we used a perplexity of 45. For the eye-only sample t-SNE, we used a perplexity of 35. The t-SNE coordinates were clustered by DBSCAN with the eps parameter set to 1.3. The cluster assignments from DBSCAN were then aggregated to the tissue and origin level, to identify small numbers of samples that clustered with other tissues; these are likely sample swaps. These outliers were removed. The script for this process is ‘outlier_identification.Rmd.’

### Differential gene expression analysis with pair-wise testing

A synthetic pan-human gene expression set was created by randomly sampling 8 tissues from each of the 22 GTEx tissue samples. This was used with the nine different eye tissue-origin sample sets and the ESC set, totaling 11 different groups. All 55 pair-wise tests (11 choose 2 equals 55) were done with the limma package with voom library size normalization, using the quantile-normalized TPM values as the input (23, 24). The script ‘differential_expression.Rmd’ contains the code for these steps.

### GO, HPO, and STRING enrichment

For GO enrichment, the biomaRt package was used, in R, to get the entrez IDs from the ‘dec2016’ ‘hsapiens_gene_ensembl’ mart. The GOstats package, in R, was used to calculate GO enrichment by the hypergeometric test, only keeping over-enriched terms. The background gene list across the different tests was defined as all genes in the original TPM expression matrix. The function for this analysis is provided as ‘GO_enrichment.R.’

For HPO enrichment, no working R package was available. To identify modules that mapped to a higher than expected number of HPO terms we used bootstrapping, comparing the number of HPO terms mapped to a module (proportional to its size) against a bootstrap distribution of the same metric. To analyze overabundance of HPO terms in a module we used hypergeometric testing, comparing the number of HPO terms in a module against the background of all genes and their associated HPO terms. The ‘ALL_SOURCES_FREQUENT_FEATURES_genes_to_phenotype.txt’ file from ‘Build #124’ was downloaded on April 4^th^, 2017 from http://compbio.charite.de/jenkins/job/hpo.annotations.monthly/lastStableBuild/. This file links gene names to HPO terms. The script that did the hypergeometric testing is provided as ‘HPO_enrichment_function.R.’

STRING enrichment p-values were computed with the STRINGdb R package. We placed all genes in each module, up to 400 (the max input possible for STRINGdb). For modules with more than 400 genes (7 retina modules and 10 RPE modules), we used the 400 genes in the module with the highest kWithin connectivity. The script for this is ‘stringDB.R.’

### Tissue-level gene block analysis with KMeans clustering and gene ontology enrichment

The differential gene expression patterns across the 55 pair-wise tests were grouped into twenty clusters, each holding groups of genes with shared expression patterns. The grouping was done with the *k*-means algorithm, in R, with 10,000 iterations and the ‘MacQueen’ algorithm. The cluster assignments for each gene was joined with the eye-tissue TPM values for the gene. The TPM values were averaged for each eye tissue, then the overall gene expression in each cluster was averaged. The TPM values, averaged by tissue, then cluster, were plotted in a heatmap. The code for this analysis is in ‘kmeans_de_cluster_heatmap.Rmd’ and the cluster assignments for each gene are available as ‘DE_Kmeans_cluster_Gene_Lists.zip’.

### Gene network construction with WGCNA

Weighted co-expression networks were constructed separately on both retina and RPE samples using the Weighted Gene Co-Expression Network Analysis (WGCNA) framework with the corresponding *WGCNA* R package. TPM expression matrices were used for the construction of both networks. Genes with consistently low levels of expression (less than 30 TPM in at least 5% of samples for the retina network, less than 40 TPM in at least 5% of samples for the RPE network) were removed prior to network construction. We found that less stringent cut-offs for low expression resulted in poor clustering of these genes (data not shown).

Average-linkage hierarchical clustering and t-distributed Stochastic Neighbor Embedding (t-SNE) were used to assess batch issues stemming from sample origin and study source, using the *WGCNA* and *Rtsne* R packages, respectively. Following the observation of batch effects, the *ComBat* R package was used to correct for batch issues stemming from an interaction variable between sample origin and study source. Following batch correction, a log_2_-transformation was applied to each expression matrix. the following transformation was applied to each expression matrix:

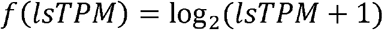

WGCNA identifies co-expression patterns using a weighted correlation matrix. The un-weighted correlation matrix is raised to a soft-thresholding power (*β*) in order to satisfy the scale-free law (25). This means that *p*(*i*), the probability that a node has degree *i*, follows a power law distribution *p*(*i*)~*i*^−*n*^. In choosing *β* for each of the networks, it is suggested by the WGCNA developers to choose a which produces a negative correlation between and log(*p*(0), with *R*^2^ > 0.8. Using the *pick Soft Threshold* function in the *WGCNA* R package, a range of soft-thresholding powers (*β*) were evaluated for both networks. The suggested criteria were met with soft-thresholding powers of 4 and 7 for the retina and RPE networks, respectively. Each co-expression network was constructed in the following manner using the-transformed expression matrices:

1. Compute a Pearson correlation matrix of all gene pairs: *S* = [*S_ij_*], where *S_ij_* = |*cor*(*i,j*)|, where *i* and *j* are distinct genes.
2. Compute an adjacency matrix as:

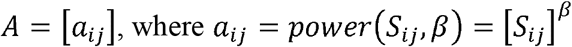
3. Compute an unsigned topological overlap matrix (TOM) as:

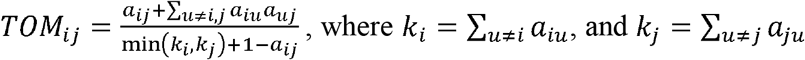
4. Define a dissimilarity matrix as *d_ij_* = 1 – *TOM_ij_*. Use average-linkage hierarchical clustering on the dissimilarity matrix to cluster the genes.
5. Use the *cutreeDynamic* function to place genes into distinct modules. For this function, parameters of *deepSplit = 0* and *minClusterSize = 30* were used.

The script used to generate the networks is provided as ‘WGCNA_networks.Rmd.’

### Identifying Similar Modules Across Retina and RPE Networks

Similarities in module compositions between the retina and RPE networks were assessed. This was performed through pair-wise cross-network comparison of retina and RPE modules in terms of the genes that were assigned to each pair of modules, as well as the GO terms that were associated with the modules being compared. For each cross-network module comparison, the number of overlapping genes was calculated and subjected to a hypergeometric test to assess significance. This process was repeated with examining overlap in GO terms between modules. In both analyses, p-values were adjusted using the FDR correction method.

### Processing of Macosko et al. single cell RNA-seq dataset

The counts from study GSE63472 were downloaded as file GSE63472_P14Retina_merged_digital_expression.txt.gz and processed with Seurat (2.3) (26). The full script is made available as process_macosko.R. Outlier cells were filtered out by removing ones which had more than 6000 or fewer than 900 genes expressed. This reduced the number of cells from 44,098 to 10,831 cells. The expression values were normalized with ‘LogNormalize’ and scaled against mitochondrial percentage and nUMI with the negative-binomial regression. t-SNE dimensionality reduction was done to visualize the expression data. The cluster assignments for each cell from Macosko et al. was downloaded from http://mccarrolllab.com/wp-content/uploads/2015/05/retina_clusteridentities.txt.

### Comparison of scRNA-seq with bulk RNA

The lists of genes in each retina WGCNA network module were pulled and their cell-type specific expression in the Macosko et al. mouse retina single cell dataset was calculated (27). High variance expression was identified by setting retina network – cell type expression with variance > 9. A fake dataset was created by randomizing the assignment of cells to retina network clusters. This fake set was used as the control group for the wilcox t test to determine whether a retina network – cell type group expression was significantly different. The code for this analysis is made available in scripts.zip as ‘single_cell_retina_network_comparison.Rmd.’

### Selection of candidate functional RPE genes and differentiation of human induced pluripotent stem cells (iPSC) into RPE cells

We took genes which: 1. were much more highly expressed in fetal RPE and stem cell RPE relative to the synthetic body set, 2. were more highly expressed in fetal RPE relative to the retina and adult RPE-choroid, and 3. had an RPE network kWithin score > 10. This produced a list of 36 genes (Supplemental Materials, Table S2). 17 of these genes were randomly selected and we also added SLC13A3 and GDF11 as these genes were common RPE network partners of the short list. Best1, MITF-Pan, MITF-M, and MITF-A were included as positives controls.

To calculate the hypergeometic p value, we counted the number of genes which were abs(0.5 log2 Fold Change) different in RPE stem cell derived relative to RPE fetal tissue (6096 out of 19128 total genes). The phyper function in R was used as 1 - phyper(14, 6906, 19128-6906, 19).

Tyr-GFP 3D1, an RPE-specific reporter hiPSC line was grown in complete Essential 8™ Medium (Life Technologies, cat# A1517001) on vitronectin (Life Technologies, cat# A14700) coated tissue culture plates at 37°C in a humidified atmosphere of 5% CO_2_. Differentiation of Tyr-GFP 3D1 cells into RPE was performed according to previously described protocols (28, 29). Differentiated RPE cells were maintained in RPE medium: MEM alpha (Life Technologies; cat # 12571-063) with 5% FBS (Hyclone; cat # SH30071-03), 1% CTS^™^ N-2 supplement (Life Technologies; cat # A13707-01), 0.1 mM MEM non-essential amino acid solution (Life Technologies; cat # 11140),1 mM Sodium Pyruvate (Life Technologies; cat # 11360-070), 250 ug/mL Taurine (Sigma; cat # T4571), 20 ug/L Hydrocortisone (Sigma; cat # H6909), 0.013 ug/L 3,3’,5-Triiodo-L-thyronine sodium salt (Sigma; cat # T5516).

### RNA isolation and TaqMan^®^ real-time PCR

GFP-positive and GFP-negative RPE cells were sorted using FACSDiva 8.0.1 cell sorter (BD Bioscience) and lysed with TRIzol reagent (Thermo Fisher Scientific; cat # 15596026). Total RNA was isolated using the Direct-zol^™^ RNA Miniprep Kit (Zymo Research, Irvine, CA) and one-microgram of total RNA was reverse-transcribed using High Capacity cDNA Reverse Transcription Kit (Applied Biosystem). TaqMan probe/primer set for the target genes were designed and gene expression was performed on the cDNA using TaqMan^®^ Universal PCR Master Mix on StepOne Plus Real-Time PCR instrument (Thermo Fisher Scientific/Applied Biosystems) (Supplementary Material, Table S3).

### Web app, other tools, and source code

The fastq file transfer and salmon quantification were run in the bash environment. The salmon-based RNA-seq quantification used the computational resources of the NIH HPC Biowulf cluster (http://hpc.nih.gov).

All other statistical analyses and visualization was done in the R environment (see ‘session_info_R.txt’ for packages used and versions). The heatmaps were made with the superheat package. All other figures were made with ggplot2.

The interactive web application was built with the R Shiny framework and hosted on a R Shiny Server (https://shiny.rstudio.com) installation at NEI. ggiraph was used to turn ggplot images into interactive images. The visNetwork R package was used to visualize the network modules. For the purpose of limiting the number of edges to a number that would be tractable for interactive visualization, the network edges were filtered so that each node would have its - nearest within-module genes (*k*-strongest edges to genes in the same module) remain in the network, for a range of *k* values.

The source code and links to the data for the web application is available at https://gitlab.com/davemcg/Human_eyeIntegration_App. The scripts mentioned in the methods underlying the data processing and analysis for this paper are available as supplemental file scripts.zip and the data used in the scripts is available at Zenodo (10.5281/zenodo.569870).

## Results

### Hundreds of individual human eye tissue RNA-seq datasets publicly available across twenty-one research studies

To identify all publicly available human eye tissue RNA-seq datasets, the Sequence Read Archive (SRA) was queried on January 19^th^ 2017 with the R package SRAdb for human transcriptomic studies with the keywords ‘RPE’, ‘macula’, ‘fovea’, ‘retina’, ‘choroid’, ‘sclera’, ‘iris’, ‘lens’, ‘cornea’, and ‘eye’ across numerous fields in the SRA (30). This inclusive search identified 603 samples across 53 studies. Hand searching the studies to identify human eye tissue samples that did not have chemical, pharmacological, or genetic modifications or known eye-disease pared the initial search down to 219 samples across 21 studies (Supplementary Materials Table S1, Fig. 1A) (15, 16, 29–43). The metadata of the remaining eye samples was queried and parsed to label each sample by tissue (cornea, retina, RPE) and origin (immortalized cell line, stem cell line, fetal tissue, adult tissue) (Fig. 1B). Before gene expression quantification and quality control to remove lower quality samples we had 110 retina, 85 RPE, 28 cornea, 16 human embryonic stem cell lines (ESC), 6 lens, and 4 eyelid tissue RNA-seq data sets.

**Figure 1.**
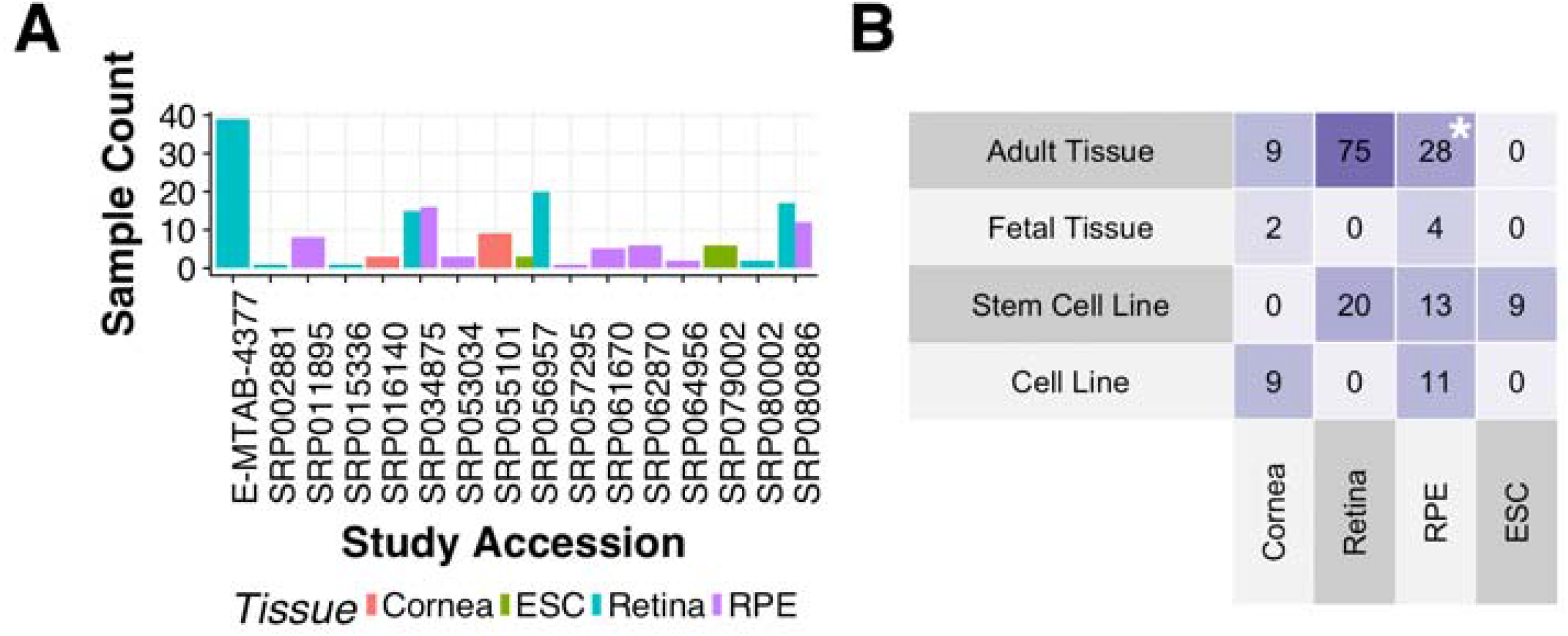
Identifying 177 unique human eye and ESC samples across 16 studies and four tissue types A. Counts for unique cornea, ESC, retina, and RPE (choroid) human RNA-seq samples by study accession B. Counts by tissue and origin. * is adult RPE – choroid

### Efficient quantification tools allow for comparison of the eye transcriptome meta-set with dozens of other human tissues

The raw sequence data was obtained from the SRA or European Nucleotide Archive (ENA) and the transcript counts were quantified with the Salmon pseudo-alignment transcript quantification (17). To improve reliability of quantification, the transcript level counts were merged to the gene level (20). We then applied quantile normalization of the TPM (transcripts per million) values on a per-tissue basis with the qsmooth tool to reduce variability between different studies (22). Outliers with extremely low median gene counts and individual samples that clustered very far apart from similar samples were removed, leaving 171 eye samples (Fig. 1A, Supplementary Material, Tables S4 and S5). Voom normalization was then applied to adjust for different library sequencing depths (23). See methods for further details.

This efficient bioinformatic process also enabled us to bring in 878 samples from the GTEx project to compare to our eye meta-set (9). We selected, when possible, 10 male and 10 female non-gender specific tissues from the GTEx, ending up with 22 tissues, including blood, brain, heart, kidney, liver, lung, and thyroid (Supplementary Material, Tables S1 and S5). All raw data from the collated eye tissues or GTEx were processed identically with the above workflow. After outlier removal, using the same workflow as the eye tissue set above, we have 853 GTEx samples across 22 tissues.

### Eye tissues from disparate studies cluster according to labelled eye component and tissue or cell-line origin

Our first question was whether the collated eye tissues, which potentially have significant batch effects from merging data from disparate sources, would group together using dimensionality reduction approaches. We used the Barnes-Hut implementation of the t-Distributed Stochastic Neighbor Embedding (t-SNE), which has been shown to work well in single-cell RNA-seq study analyses as well as the GTEx study set, to visualize relationships in two dimensions between the processed eye tissues (Fig. 2A) (21, 46, 47). The DBSCAN algorithm was used on the t-SNE coordinates for each sample to identify nine distinct clusters (48).

**Figure 2.**
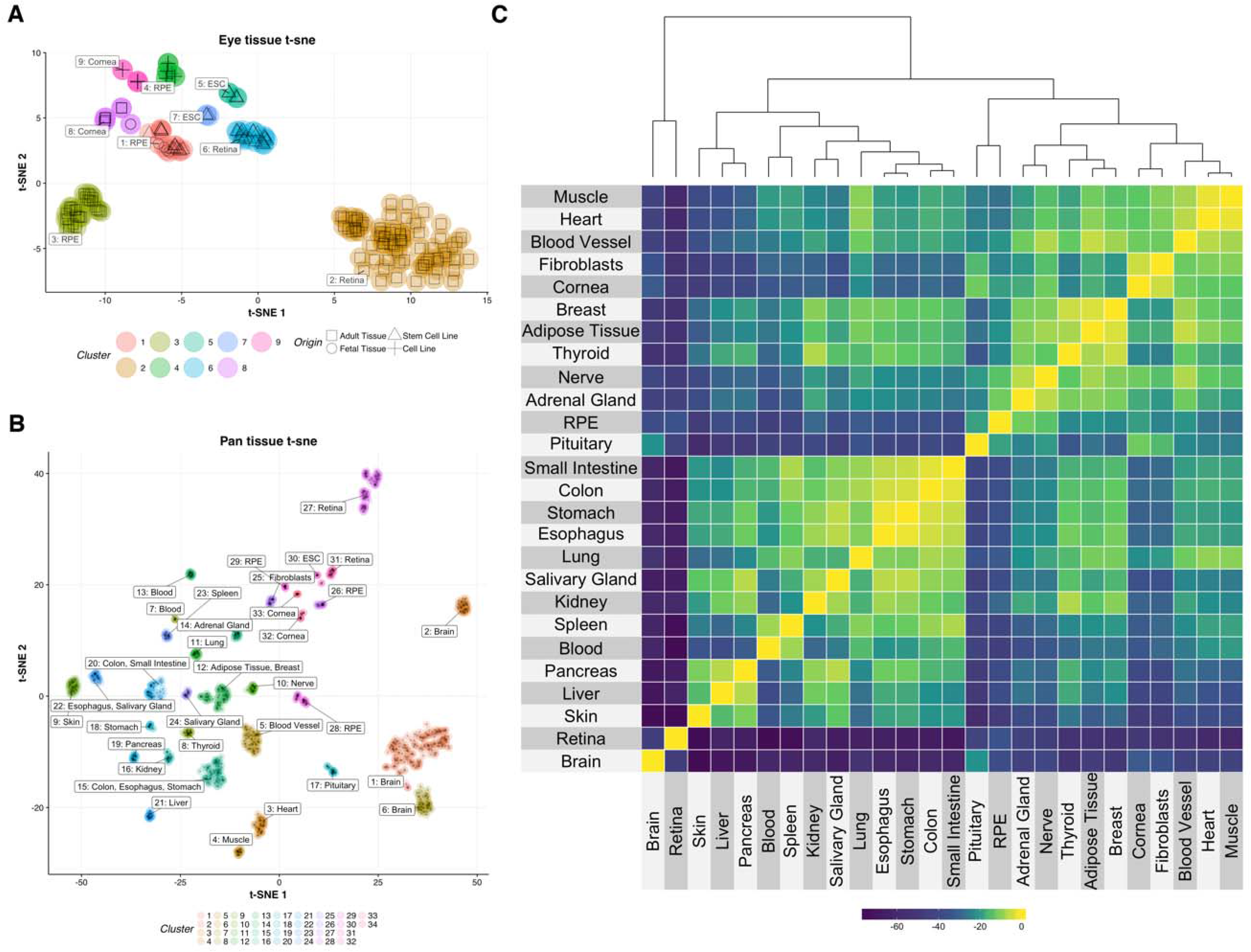
Gene expression information sufficient to both accurately cluster eye and GTEx tissue independently and demonstrates that eye tissues are generally more closely related to other than other body tissues A. Dimensionality reduction by t-SNE of human eye tissues and ESC, colored by clustering assignment, and labelled by tissues in cluster. Shape of point corresponds to tissue origin B. Eye tissues with GTEx tissues, colored by clustering assignment, labelled by tissues in cluster C. Pair-wise euclidean distance between each tissue. Closer tissues have a smaller height in the dendrogram and are more yellow in color. More distant tissues have a larger height in the dendrogram and are more blue.

The adult tissue retina samples clustered together, though apart from their fetal or cell line based samples. The ESC-derived retina samples have a variety of time points (37, 47, 67, 90 days) during their differentiation; we found no clustering by those criteria (data not shown) (33). The fetal and adult cornea samples, grouped closely together, but still clustered independently (Fig. 2A, clusters 8 and 9). Human embryonic stem cells (ESC), included because they are used across several studies to differentiate into different eye tissues, clustered together, generally closer to the cell-line derived samples (Fig. 2A, cluster 5).

RPE is the only tissue for which more than three different sources were available: fetal tissue, adult tissue immortalized cell-line, and cells differentiated from ESCs. It should be noted that the adult RPE tissues are a mixture of RPE and choroid tissue, which is a vascular layer of the eye, providing oxygen and nutrients to the RPE and outer retina. This tissue will be referred to as adult RPE/choroid. The four sources cluster into three groups, with the few RPE fetal tissues clustering with the ESC-derived RPE samples (Fig 2A, cluster 1). The RPE derived from ESC group (Fig 2A, cluster 1) is composed of samples from three studies (31, 36, 41). All three groups differentiated their RPE cells for about two to four months, according to their method section. Wu and Zeng et al. gave specific times of differentiation for the exact tissues used in the SRA metadata (40 or 100 days); we did not see any differences in clustering patterns based on length of differentiation (data not shown) (41). This close grouping of fetal and ESC-derived RPE tissues are consistent across multiple runs of t-SNE with different perplexity parameters ranging from 35-50 (data not shown). The adult RPE/choroid tissue clusters further away from the cell-line based tissues.

Overall, the t-SNE dimensionality reduction demonstrates that the eye tissues consistently cluster in unique groups by their tissue and origin. This happens despite a variety of laboratory origins with disparate culturing conditions, tissue handling, RNA extraction, sequencing cores, and so on.

### Eye tissues distinct from most human tissues

To explore the relationship of eye tissues to other tissues in the human body, we leveraged the GTEx data we reprocessed to create a pan-human two-dimensional tissue relationship map with t-SNE (Fig. 2B). DBSCAN was then used, as before, to identify clusters. ‘Tissue’ labels from GTEx metadata in the SRA were used with one exception; fibroblasts are labelled separately from ‘Skin’ as they consistently group independently of skin-punch tissues. From the t-SNE visualization (Fig. 2B) we observe most human tissues group close to each other, with the exception of brain. The eye tissues, except retina, group closer to the non-brain human tissues. While the cell-line versus tissue derived eye tissue distinctions are maintained with the pan-human set, the eye-tissues are generally more related to each other than non-eye tissues.

The t-SNE 1 and 2 dimension coordinates generated by t-SNE are sensitive to the parameter perplexity, which controls the weighing of local to global relationships (49). Figures 2A and 2B used perplexities of 35 and 45, respectively. To more consistently demonstrate the pair-wise relationships between the tissues, the t-SNE dimensions were iteratively generated with perplexities from 35 to 50. Then means were taken, grouped by sample. The individual samples were then grouped by labelled tissue type and the t-SNE coordinates were again averaged. Hierarchical clustering by Euclidean distance was done to group the tissues and a heatmap was generated (Fig. 2C) which displays the most closely related tissues. Because the hierarchical distances between cell-line derived eye tissues were inconsistent, they were removed from this analysis. We see that retina and brain tissues are individual outliers. We also see that the pituitary is grouped near RPE tissue and that fibroblasts group closely with the cornea (as denoted by the height of the dendrogram).

### Differential expression analysis identifies large sets of genes distinguishing separating eye tissues

The eye tissue set collected can be separated on two major axes: tissue type (cornea, retina, or RPE) and origin (immortalized cell line, stem cell line, fetal tissue, adult tissue). Labelling each set of tissues by these two criteria gives us ten sets of eye tissues (Fig. 1B). To compare expression against non-eye tissue, we created a synthetic human ‘body’ expression set, by evenly combining the 22 GTEx tissues. The total number of body samples was matched to the total number of eye tissues we have by taking a random set of 8 tissues from each human body tissue category (e.g. Brain, Pituitary). There are 55 two-way combinations possible among the 11 sets.

To calculate differential expression, we modeled expression with the limma linear fit function with voom to correct for library size differences. The limma empirical Bayes function was used to identify statistically significant differentially expressed genes (23, 24). To look for global changes between the eye tissues and the body, we will first compare all of the eye tissue groups individually against the synthetic body (Table 1). A second synthetic body set was created by sampling the un-used GTEx tissues from the first synthetic body set and we found very similar differential expression values (data not shown).

**Table 1.**

| Comparison | Log Fold Change |  |
| --- | --- | --- |
|  | < -2 | > 2 |
| Cornea (adult) vs Body (adult) | 1969 | 1249 |
| Cornea (fetal) vs Body (adult) | 172 | 873 |
| Cornea (immortalized cell) vs Body (adult) | 2611 | 1475 |
| ESC (stem cell) vs Body (adult) | 2738 | 2177 |
| Retina (adult) vs Body (adult) | 2607 | 1978 |
| Retina (stem cell) vs Body (adult) | 3443 | 2622 |
| RPE - choroid (adult) vs Body (adult) | 1200 | 1258 |
| RPE (fetal) vs Body (adult) | 1510 | 1402 |
| RPE (immortalized cell) vs Body (adult) | 2446 | 1398 |
| RPE (stem cell) vs Body (adult) | 2270 | 1308 |

Many genes are differentially expressed between the eye tissues and the synthetic body
Number of genes with logFC < -2 or > 2 (0.25 or 4 fold, p value < 0.01) between each eye tissue against the synthetic body set

The differentially expressed genes identified for each test (Table 1) was filtered to retain only genes with log fold change (logFC) < −2 or > 2 relative to the baseline tissue and with a false discovery rate (FDR) corrected p-value less than 0.01. A logFC of more than two means that the detected transcript level is more than four times as much (or one quarter as much) compared to the body tissue.

### Biological term enrichment identifies eye-specific gene expression biology relating to visual function and body-specific gene expression relating to immunity and cell adhesion

As we have hundreds to thousands of genes meeting these stringent differential expression criteria across the ten comparisons we did Gene Ontology (GO) biological process term enrichment to identify systems-level patterns. We did the GO term enrichment independently on the over- and under-expressed gene sets, relative to the synthetic body set; 20 tests were performed. Overall, we found 2796 unique GO term IDs across the tests with an FDR corrected p value under 0.01 (Supplementary Materials, Table S6).

We took the top forty GO term IDs from the over and under-expressed tests (ranked by p value) and plotted them in a heatmap to identify shared GO terms among the different comparisons and to find overall trends in eye tissues gene expression relative to the synthetic body gene expression set (Figure 3). Clustering was done on both rows and columns to group together shared patterns. Like the t-SNE based clustering, the retina is an outlier for GO term enrichment. The GO terms in the first 20 rows (Fig. 3, Block 1) is driven by genes that are more highly expressed in the retina relative to other tissues. These over-expressed genes are highly enriched in GO terms relative to visual perception, light stimulus, synaptic signaling, and neurogenesis.

**Figure 3.**
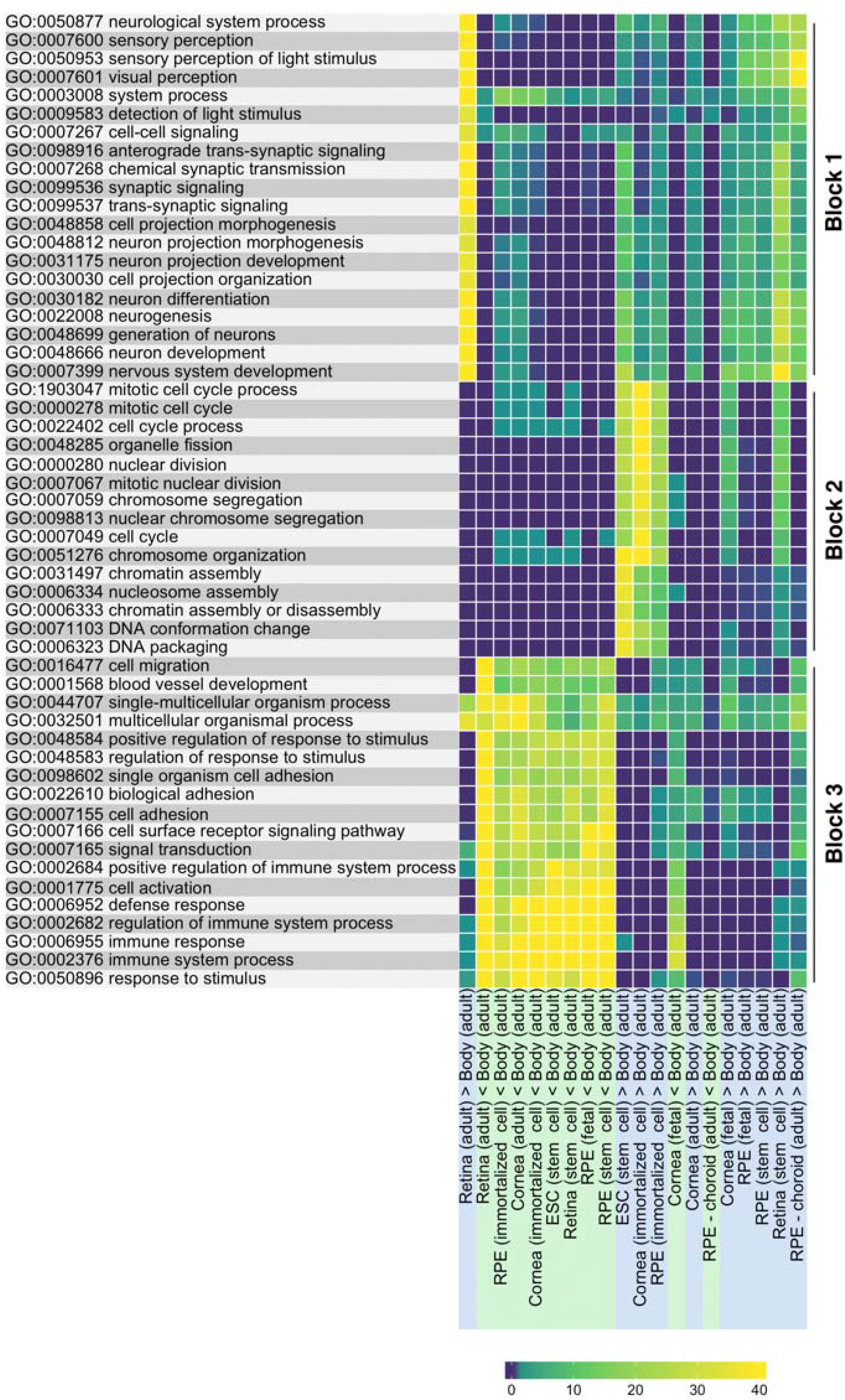
Major differences in systems relating to visual function, active cell division, adhesion, and immunity between the eye tissues and the other tissues in the human body Top 80 GO Terms (40 with eye > body and 40 with body > eye) across eye-tissue to body differential expression tests. Yellow is more significant, blue is less. Hierarchical clustering of both rows and columns place more related GO terms and tissue comparison sets together.

The next group (Fig. 3, Block 2) of enriched GO terms most strongly defines the ESC, the cornea, and RPE immortalized cell lines and to a lesser extent, fetal cornea and stem cell retina tissue. These GO terms relate to cell cycle and division as well as DNA packaging and conformation. The last block (Fig 3., Block 3) is a set of GO IDs related to the body gene expression being higher than most of the eye tissues. This large block has GO terms involving migration, organismal process, adhesion, immune process, and stimulus. The full set of significantly (p < 0.01) enriched GO terms (2796) is available in Supplementary Materials Table S6.

### Within eye tissue differential expression comparisons identify cornea, retina, RPE, and RPE/choroid gene sets

To more directly identify sets of genes enriched in particular eye tissue(s) relative to the other eye tissues, we compared all eye tissue differential expression pair-wise against each other and the synthetic body set (55 tests). To identify common gene sets, we used *k*-means clustering to group all genes into twenty groups; each group has a different overall gene expression pattern. We then plotted the relative gene expression for each eye tissue across the twenty *k*-means groups (Supplementary Materials, Figure S1). This produces a heatmap which identifies sets of genes that are more highly (or lowly) expressed in particular eye tissue(s) relative to the other eye tissues. We use this heatmap to identify genes defining the cornea, retina, RPE, and adult RPE/choroid and did GO term enrichment on these clusters (Table 2, Supplementary Table S7). The gene lists for each of the 20 groups are available in Supplementary File S1).

**Table 2.**
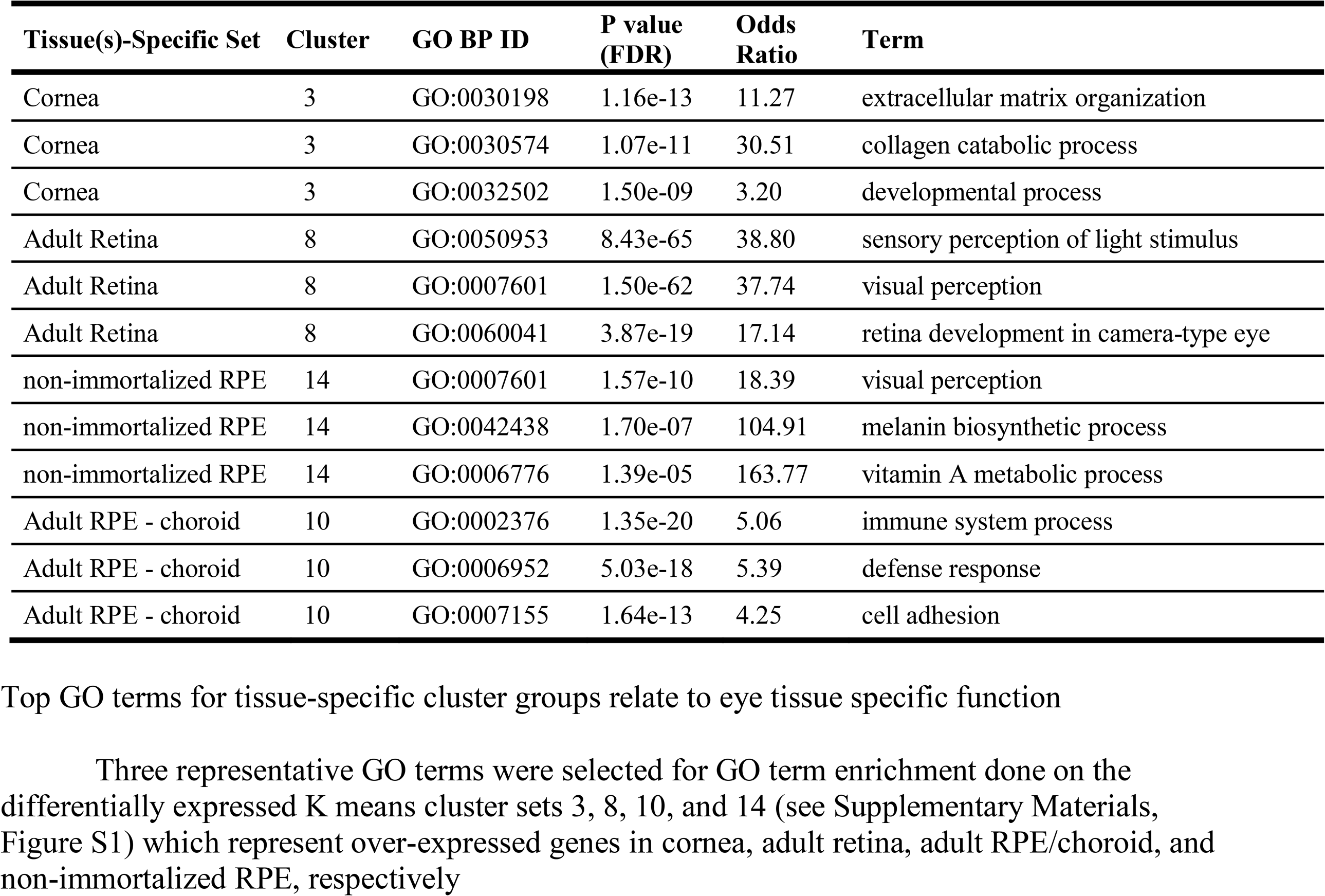

### The cornea is enriched for genes involved in the extracellular matrix and collagen relative to the other eye tissues

In the GO heatmap (Fig. 3) the cornea tissues (immortalized cell line, fetal, adult) lack a highly distinguishing set of GO terms from the other eye tissues. However, there is a cluster (Supplementary Materials, Figure S1, cluster 3), with enriched fetal and adult cornea expression compared to the other tissues. This cluster contains 157 genes and top GO terms enriched for this set relate to extracellular matrix organization, collagen metabolism, and developmental processes (Table 2, Supplementary Table S7).

### Adult retina and, to a lesser extent, retina stem cells enriched in visual function genes

Compared to the synthetic body set, the adult retina has many GO terms relating to visual function (Fig. 3, Block 1). This same GO enrichment is seen even when comparing adult retina against the other eye tissues, focusing on cluster 8 (Supplementary Materials, Figure S1, Table 2). This cluster is very highly expressed in adult retina and somewhat highly expressed in stem cell derived retina, relative to the other eye tissues.

### RPE, excluding hTERT RPE, is highly enriched in genes relating to pigmentation and visual perception

Like cornea, the non-immortalized RPE tissues do not have a distinct block of GO terms (Fig. 3). In the *k*-means heatmap (Supplementary Materials, Figure S1) we see that cluster 14 is more highly expressed in stem cell RPE, fetal RPE, and adult RPE/choroid. The hTERT immortalized cell line RPE is not highly expressed for this gene set. The 92 genes in this cluster are enriched in GO terms for visual perception, melanin processing, and vitamin A metabolism (Table 2).

### Compared to other eye tissues, adult RPE/choroid is enriched for genes involved in immune function and adhesion

The cluster with genes highly expressed in adult RPE/choroid compared to the other eye tissues (number 10), has 229 genes. As this cluster is not highly expressed in the other RPE tissues, this cluster may define the choroid. These genes are strongly enriched in immune function and adhesion (Table 2).

### hTERT RPE immortalized cell line has substantial gene expression differences relative to RPE derived from ESCs

As we had seen that the hTERT RPE clusters apart from the other RPE tissues, and there is a benefit to examining the differences between an immortalized RPE cell line model versus a differentiated RPE cell line model, we looked directly at differences in expression between hTERT RPE and stem cell derived RPE. We identified what genes and GO terms make these two cell lines different. There are over 1323 genes with a more than four-fold expression difference between RPE derived from human ESCs and the ATCC hTERT RPE immortalized cell line and 1572 with four-fold lower expression (Supplementary Materials, Table S8). The five genes most highly expressed in RPE derived from human ESCs relative to the ATCC hTERT RPE immortalized cell line are *TTR* (Transthyretin), *DCT* (Dopachrome Tautomerase), *KIF1A* (Kinesin Family Member 1A), *SFRP5* (Secreted Frizzled Related Protein 5), and *NELL2* (Neural EGFL Like 2). GO terms associated with higher stem cell RPE expression relate to ion transport and synaptic transmission, suggesting that stem cell derived RPE is a more faithful model of human biology (Fig. 4).

**Figure 4.**
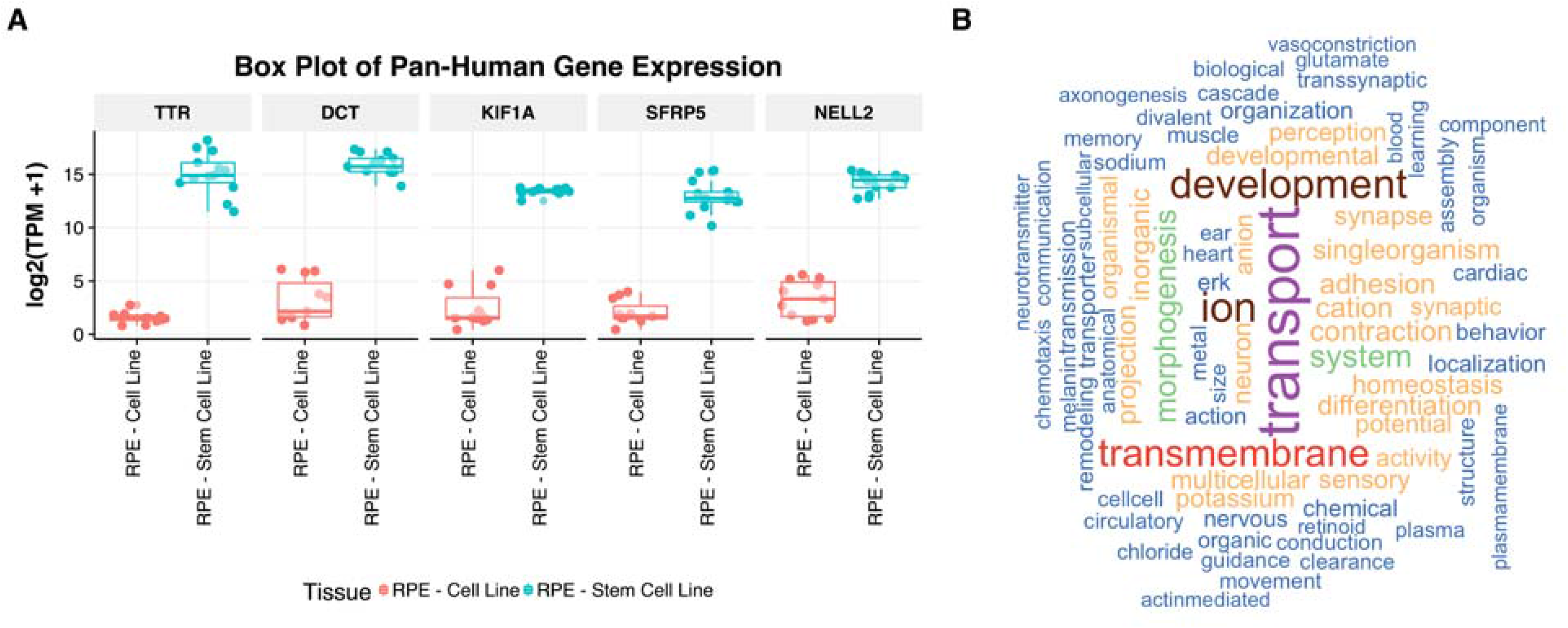
Genes crucial in eye function are highly differentially expressed between stem cell derived RPE and hTERT RPE A. The top 5 genes overexpressed in ESC-derived RPE and immortalized cell line hTERT RPE B. Word cloud of enriched GO terms

### Dissection of high expression retina genes with single cell RNA-seq reveals blocks of genes with retina-cell specific function and candidate signature genes

We took advantage of the availability of a retina single cell RNA-seq data set from normal mouse retina (P14) from Macosko et al. (27). The raw counts from 44,098 individual dissociated retina cells were filtered down to 10,831 high quality cells, reanalyzed (see methods) and clustered with t-SNE (Supplementary Materials, Figure S2). The gene expression was grouped by the eleven major cell types identified by Macosko et al. and the expression was split into deciles of expression, with 10 being genes in the top 10% of expression for the cell type. This 179, 112 row data table is made available on eyeintegration.nei.nih.gov (Data Table -> Mouse Retina Single Cell RNA-seq Data).

This expression set was combined with the list of genes expressed highly in retina (adult) relative to the synthetic body set and this list was further subdivided by whether the gene was highly expressed in any specific retina cell type. Of the 11,660 genes with > 1 fold change in the adult retina tissue over the synthetic set 1,913 were expressed in the top decile in one of the 11 retina sub-types. We ran a bootstrap test 10,000 times to calculate which of the 11 cell types were enriched in the 11,660 gene set, relative to a random set of genes of the same size. We found that amacrine, bipolar, cones, rods, horizontal, Müller glia, and retinal ganglion cells were enriched (p < 0.05).

To leverage this more specific gene list to identify functional modules we took the genes in the top decile expression for each cell type and ran GO enrichment (Supplementary Materials, Figure SX3). GO terms relating to visual perception and light stimulus are highly enriched in rods and cones and enriched in amacrine, bipolar, astrocytes, microglia, and Müller glia. Retinal ganglion cells are enriched in a set of GO processes describing neuron projections, axogenesis, and microtubule processes. Retinal ganglion, amacrine, and bipolar cells share a set of GO terms involved with synaptic vesicles, ion regulation, and neurogenesis.

To identify candidate signature genes for each cell type we looked for genes overexpressed in the bulk retina tissue relative to the synthetic body set, in the top 20% of expression in a particular retina cell type, and in the bottom 50% for the remaining retina cell types (Supplemental Table S9). Several examples of these genes are plotted in t-SNE plot of the clustered single cell retina data, demonstrating how genes like GAD1, PDE6C, and TUBB2B distinguish amacrine, cones, and retinal ganglion cells from the other retina types (Supplementary Materials, Figure S4).

### Highly connected genes in retina and RPE gene networks recapitulate known eye biology

To this point, we have used the full gene expression set to independently cluster samples by tissue type and origin. We then used differential expression between the eye tissues and the synthetic body set to highlight differences in GO terms. We delved further by clustering the differential expression patterns between the eye tissues to find how each eye tissue is different from the other. We can go even further, by examining the relationships of the genes to each other, within a tissue, by using gene correlation networks. These networks use correlated fluctuations of all-by-all pairwise gene expression similarities to build networks of gene-to-gene relationships.

As we had collected a substantial amount of retina and RPE samples, we were able to build weighted gene correlation networks with the Weighted Gene Co-Expression Network Analysis (WGCNA) R tool (25). We also attempted to build a cornea network, but the network construction failed due to failure to both differentiate the genes cleanly into defined modules and achieve appropriate network topology within a reasonable parameter space; more cornea samples are needed (Supplementary Materials, Figure S5). The gene expression TPM values, with the full set of corrections described earlier for the differential expression analyses, were used as inputs. All retina and all RPE tissues that passed quality control steps were used to build independent retina and RPE networks. The parameters used in the WGCNA network construction are enumerated in the methods.

There are 11,101 and 10,843 genes in the retina and RPE networks, respectively. 9621 of the genes are shared between the retina and RPE network. The kWithin metric from WGCNA measures the intramodular connectivity. Genes with higher connectivity are, theoretically, more likely to be important in gene regulation as perturbations in them will affect the system more than less connected genes.

To get a sense of what the biology was of the most connected genes in the retina network, we took the 1017 genes with a kWithin greater than 20 and did GO enrichment (Supplementary Materials, Table S9), finding the top five GO terms all relate to visual perception. We did the same with the RPE network, using the 566 genes with a kWithin greater than 20. The top five GO terms in this RPE network connected list were related to endoplasmic reticulum function (Supplementary Materials, Table S10). The most similar modules, calculated by doing hypergeometric testing of GO terms and gene names, between the retina and RPE networks are the light cyan retina module and the pink RPE module. Both of these modules, by GO term enrichment, are involved in protein targeting to the ER (Supplementary Materials, Figure S6).

### Retina network module highly enriched in genes implicated in eye disease and crucial for visual function

A key advantage of WGCNA networks over correlation networks is that genes can be partitioned into modules, presumably with shared biological function within each individual module. The retina network has 27 modules, with 64 to 1922 genes in each module. The RPE network has 23 modules, with 90 to 1458 genes in each module (Supplementary Materials, Figure S7). To determine whether the modules were enriched for known gene to gene interactions, we loaded each network module gene list into STRING and calculated whether there were more interactions than expected. For 23/27 retina modules and 20/23 RPE modules, the STRING p value for interaction enrichment was < 0.01 (Supplementary Materials, Table S10). We also ran GO term enrichment for each module within each network (Supplementary Materials, Table S14 and S17). While many modules have highly significant GO term enrichment, only the ‘green’ module is highly enriched for visual perception terms. Pinelli et al. built an unweighted retina gene correlation network and identified 14 candidate photoreceptor genes based upon their network (16). All 14 are in our retina network and 9 of the 14 are in our green visual function module (p < 2.8 × 10^−10^) (Supplementary Materials, Table S12).

There are 617 genes within the green retina module and 178 of these have a kWithin greater than 20. Many of the top connected genes have known visual function or are implicated in retinal diseases. To demonstrate the strong enrichment of known eye function genes in this module we divided the genes in the green module into four categories: known to play a role in eye disease, having GO terms relating to visual function, both, or neither (Fig. 5, Supplementary Materials, Table S13). From RetNet (http://www.sph.uth.tmc.edu/RetNet/) we have a list of 331 genes that have been implicated in retinal diseases (5). There are 178 genes with kWithin > 20 in the green module; 14 of those genes are also in RetNet, 17 have a vision GO term, 31 have both, and the remaining 116 genes are neither in RetNet nor have a vision-related GO term.

**Figure 5.**
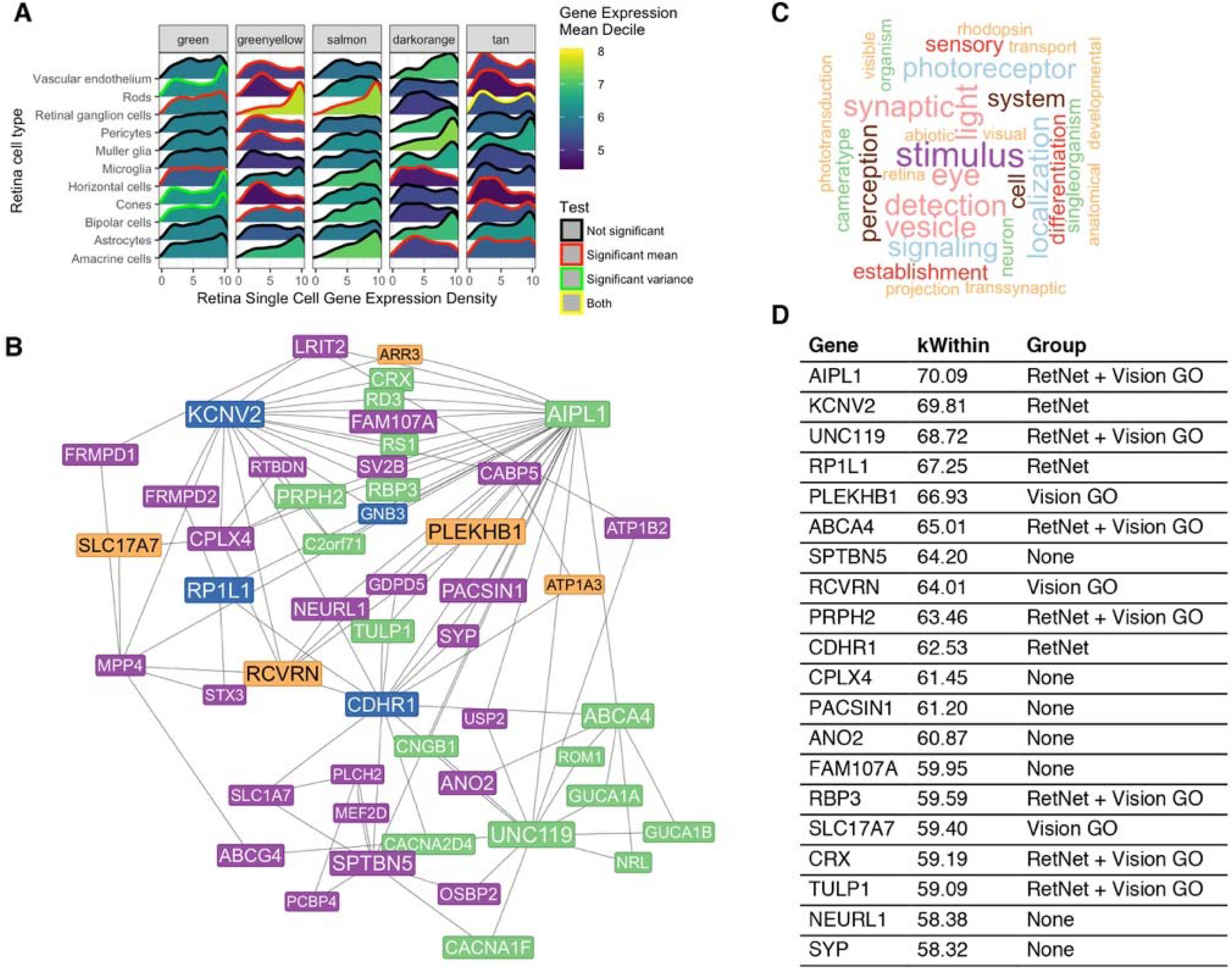
Retina network green module highly enriched for important visual function genes A. Density plot showing expression of retina network module genes across different retina cell types in single cell mouse retina RNA-seq. Group differences in mean expression and variance are shown by the line colors in the density plot. B. Top 50 connected genes in green module in the retina network. Colored by group (see C.) C. Word cloud of top GO terms in green retina module D. kWithin connectivity (higher is more connected) for top 20 connected genes, labelled to indicate whether the gene is in RetNet, has a GO term relating to visual function, both, or none.

The human phenotype ontology (HPO) project is conceptually similar to gene ontology, except that they map abnormal human phenotype terms onto a graph and match them to genes (50). This provides a way to identify enrichment of abnormal human phenotypes. As there is no functioning package in R to systematically calculate HPO enrichment, we did bootstrapping and hypergeometric testing (see methods), looking for enrichment overall at the module level and for individual HPO terms within each module, respectively. The green module is highly enriched for HPO terms relating to eye disease, with terms like nyctalopia, abnormal electroretinogram, photophobia, cone-rod dystrophies, and blindness among the top terms (Supplementary Materials, Table S15).

Other highly significant GO terms in the remaining retina network modules also match known retina function. GO terms enriched relate to ion transport (greenyellow), developmental processes (darkorange, greenyellow, tan), mitochondrial function (midnight blue), and metabolism (turquoise) (Supplementary Materials, Table S14). The retina network darkgrey module also contains several genes implicated in retina diseases like *ELOVL4, OPN1SW, SLC24A1*, and *PDE6A* (see Supplementary Materials, Table S16 for full list). Additionally, the green, tan, brown, and blue modules are, overall, enriched for HPO disease terms (Supplementary Materials, Figure S8).

### Differentially expressed, high connectivity RPE genes are highly expressed in functional pure RPE cells

To experimentally validate whether our differential RPE gene expression data and RPE-choroid network connectivity could identify important genes in functional RPE we first made a short list of highly expressed and high connectivity RPE genes (see methods). We then compared expression of the genes in human iPSC-derived RPE, purified using an RPE-specific *TYR* enhancer coupled to a GFP transgene. Differentiating cells were then sorted using flow cytometry to purify the GFP positive cells in the population. We find that 17 of our 19 genes are more highly expressed in the GFP+ RPE (Supplementary Materials, Figure S9). 14 of the 19 genes are 0.5 log2 fold change greater than the in the purified RPE population (hypergeometric p < 0.0002, see methods).

### Retina green module identifies visual transduction pathway and core upstream regulators

The green module was further analyzed for known biological networks components, which were generated through the use of Ingenuity Pathways Analysis (Ingenuity^®^ Systems, www.ingenuity.com). Visual transduction was the most significant pathway present, with 16 components present in the green module. These components function predominantly in rod and cone photoreceptors in the conversion of photic energy to neural signaling in the retina (Supplementary Materials, Figure S11A and data not shown), as confirmed by the single cell RNA-seq dataset. Regulatory component analysis projected that *CRX* and *NRL* were predicted among the regulators of gene expression in the green module, upstream of several genes implicated in retinal photoreceptor degeneration also present in the green module (Supplementary Materials, Figure S11B). These two transcription factors drive rod photoreceptor differentiation and maintenance beginning in embryogenesis, and dysfunction of either of these is associated with retinal degeneration (51). In sum, the green module is enriched for photoreceptor function and recapitulates specific components of known biological and gene regulatory networks that are important causes of retinal disease.

### RPE/choroid network contains many modules related to cell metabolism

Unlike the retina network, there are no strongly associated GO terms relating to visual function. However, there are numerous modules with strongly significant GO terms relating to metabolic processes and active transcription and translation (blue, brown, dark turquoise, green, light cyan, light green, red, turquoise). One module (yellow) relates to catabolism, one to immune function (tan), one to the endoplasmic reticulum (ER) (pink), and two the mitochondria (dark green, dark yellow) (Supplementary Materials, Table S16). Among the top HPO terms across the RPE modules are ones relating to anemia (pink), optic disc pallor (green), and respiration (dark green). Overall, the green, midnightblue, turquoise, lightyellow, magenta, and brown RPE modules are enriched for HPO terms (Supplementary Materials, Figure S9).

### Single-cell RNA-seq confirms many network modules represent specific retina cell types

Because the GO terms are distinct between the different WGCNA modules some of them should be enriched in genes distinguishing different types of retinal cells. We took the Macosko et at retina single cell sequencing data and looked at the expression profile for each labelled cell type grouped by retina network module color (Supplementary Materials, Figure S10). We see that many modules have very similar expression profiles across the twelve retina cell types (see black, blue, brown, darkgreen).

However, we also see that several modules have strongly divergent expression patterns between different cell types (Figure 5a). For example, the green module is enriched in genes with strong expression in rods, cones, and bipolar cells. The greenyellow and salmon modules have genes with high expression in retinal ganglion cells. The darkorange and tan modules are enriched for genes expressed highly in Müller glia and astrocytes.

### Retina and RPE networks in retinal diseases and AMD

Higher connected genes are theoretically more important in the function of the retina and RPE. From RetNet we have a list of 331 genes that are associated with retinal diseases (though some unknown proportion affect the retina via the RPE). From a recent large AMD GWAS study, there is a list of 33 loci strongly associated with AMD, and thus likely related to RPE or choroid dysfunction (8). To see whether these retina or RPE gene lists have higher connectivity relative to the other genes in the networks we used density plots of the kWithin value to see whether we see any left-ward (less connectivity) or right-ward (more connectivity) shifts in our gene list kWithin connectivity.

We see that the RetNet gene list has a higher connectivity than non-RetNet genes in the retina module; this right-ward shift is highly significant (p = 3.26 × 10^−8^). The connectivity of the RetNet gene list in the RPE network is significantly different than the non-RetNet genes (p = 0.28). 53 RetNet genes are in the green retina module, which is a 4.1 fold enrichment over chance. The darkgrey module has a similar enrichment in RetNet genes with 10, which is a 3.8 fold enrichment over chance.

The 33 genes associated with AMD have a higher connectivity the remaining genes in the RPE network; this right-ward shift is also significant (p = 0.049). Like the RetNet retinal disease gene list in the RPE network, the 33 AMD genes are not significantly more connected than the other genes in the retina network (p = 0.49) (Supplementary Materials, Figure S12).

## Discussion

We collected all publicly available human eye RNA-seq datasets creating the largest pan-eye collection to date, and carefully performed a lengthy series of normalization and quality control procedures to robustly quantify gene expression within three major eye tissues and between the eye and other human tissues. Gene expression data can be used to accurately cluster samples by tissue and origin. We used differential gene expression analysis with GO term enrichment to identify biological processes that best distinguish the eye tissues both from each other and from a synthetic human expression set. We then leveraged the large sets of retina and RPE tissues to build the first human weighted gene correlation networks for retina and RPE, confirming with single cell RNA-seq data that several of the retina modules have cell-type specific expression. We demonstrated the power of the networks to highlight genes known to be crucial in eye biology. Finally, we make the data and analyses available in a powerful web application (https://eyeIntegration.nei.nih.gov).

The structures of the eye are epithelial, neuroepithelial, and neural crest in origin. We were expecting some of the eye tissues to cluster closely with the skin, but instead we found that the retina was a very unique tissue, that transformed fibroblasts most closely matched the cornea, and the RPE was nearest the pituitary. Embryologic origins and specialized functions likely create these similarities and divisions, respectively. Cornea is derived from the same surface ectoderm as skin and from neural crest cells, while retina and RPE are derived from the neural tube epithelium from the ventral diencephalon, along with the hypothalamus and posterior pituitary. Corneal epithelium is replenished by limbal stem cells that remain into adulthood, which may explain the proximity of corneal and ESC clusters. This also may reflect that the majority of our corneal dataset is derived from cultured corneal epithelium. That retina was separated from other ocular and non-ocular tissues likely related to the exclusivity and high expression burden of the visual transduction cycle in cone and rod photoreceptors.

The systems-level study of differential gene expression across cornea, retina, RPE, and RPE-choroid tissue highlights core functions of these tissues. Cornea-specific genes specify the structural aspect of the cornea with extracellular matrix organization and collagen metabolism and catabolism. The corneal epithelium is replenished continuously with limbal stem cells, which may be reflected in the enrichment of GO terms relating to development. The retinal tissues are strongly defined by genes involved in visual processes. The RPE and RPE-choroid tissues are also distinguished, with the former being more involved in visual processes and pigmentation while the latter is involved with immune system processes.

The creation of the first human retina and RPE weighted gene correlation networks has allowed us to identify dozens of modules with co-regulated genes. It is important to stress that these networks were built only with gene expression information and were optimized using network-specific metrics, such as how well the topological overlap matrix placed genes into well-defined modules. Only afterwards did we evaluate the significance of connected genes and modules to GO terms and known eye biology.

It is striking that the some of the most significant GO terms, by p-value and enrichment, in the retina network are associated with a single 617-gene module underlying visual function. This module represents the visual transduction pathway, which is relatively unique to the retina and is associated with isolated and nonsyndromic retinal degenerative conditions. Single-cell RNA-seq data demonstrated that the genes in this module are expressed more highly in the rods, cones, and bipolar cells.

As the RPE has a high-energy role in transferring nutrients and clearing waste products for the photoreceptors of the retina, it is not surprising that a plurality of the modules are enriched for genes important in RNA translation, protein modification and production, catabolism, and mitochondrial function. The enrichment of highly connected AMD associated genes in the RPE network further emphasizes the value of this network.

Finally, the value of this extensive and carefully curated data-set is enhanced by the creation of the eyeIntegration web app (http://eyeIntegration.nei.nih.gov; Supplementary Materials, Figure S13). The site serves two roles, first as an interactive extension of this manuscript and second as a platform for researchers to identify interesting genes in eye function via searchable gene expression plots across many tissues, 55 pair-wise differential expression tests, and two gene networks. We also make the source code and accompanying data-sets fully and freely available for other researchers (see methods) and will periodically refresh the data. The unravelling of eye biology and function has been furthered by genetic eye diseases, animal models, and functional assays. We hope that this open data sharing and powerful web application will provide a fourth way to decipher eye biology in health and disease.

## Acknowledgements

This work utilized the computational resources of the NIH HPC Biowulf cluster (http://hpc.nih.gov). We would like to thank the 21 eye research projects whose public sharing of their raw sequencing data and metadata made this project possible.

## Abbreviations

AMD: Age-related macular degeneration
ESCs: Human Embryonic Stem Cells
GO: Gene Ontology
GTEx: Gene Tissue Expression Project
HPO: Human Phenotype Ontology
iPSC: Induced Pluripotent Stem Cell
logFC: log Fold Change
RPE: Retinal Pigment Epithelium
SRA: Sequence Read Archive
TPM: length-scaled Transcripts Per Million
t-SNE: t-Distributed Stochastic Neighbor Embedding
WGCNA: Weighted Gene Co-Expression Network Analysis

